# Neuropeptides Involved in Elicited Reversal Speed Plasticity in *C. elegans* During Mechanosensory Habituation

**DOI:** 10.1101/2025.11.24.690280

**Authors:** Audrey Siu, Nikolas Kokan, Catharine H Rankin

## Abstract

*Caenorhabditis elegans* respond to mechanosensory taps with brief reversal responses. In past research, speed of the response was averaged over the entire reversal for each tap to analyze reversal speed habituation; however, with this measure, only a modest decrement in speed was observed. Using a more detailed breakdown of reversal speed, we found that speed is most plastic early in the reversal and stable later on. Using this analysis, we found that worms with mutations in neuropeptide genes show reduced speed plasticity during the first second of reversals, indicating peptidergic signaling may be involved in reversal speed plasticity.

## Description

Habituation is a type of non-associative learning where animals display a learned reduction in response to a repeated stimulus (Rankin et al., 2009). This phenomenon is highly conserved across many species, including *Aplysia californica* (Glanzman, 2009), *Drosophila melanogaster* (Engel and Wu, 2009), zebrafish (Beppi et al., 2021), rodents (Leussis and Bolivar, 2006), and humans (Thompson, 2009). Our work focuses on the nematode *Caenorhabditis elegans* (*C. elegans*), which habituates in response to repeated mechanical stimuli (Rankin et al., 1990; Ardiel et al., 2010). When stimulated by a tap to the side of the Petri plate, *C. elegans* initiate a brief reversal response known as the tap-withdrawal response (TWR) (Rankin et al., 1990; Wicks and Rankin 1995; Wicks and Rankin 1996). With repeated taps, decreases in both the frequency and amplitude of this response are observed.

Using our high through-put machine vision tracking system, the Multi-Worm Tracker (MWT), we simultaneously recorded the behaviour of dozens of worms in response to a metal solenoid striking the side of the Petri-plate holding the worms (Swierczek et al., 2011). This allowed us to assess many different response components of the TWR including reversal probability (the proportion of worms reversing to taps; Figure 1A), and reversal duration (the length of time of the reversal for worms that responded to the tap; Figure 1B). The average speed of the tap elicited reversal has also been measured, as seen in Feresten et al. (2023), Kepler et al. (2022), Mc Diarmid et al. (2020), and Ardiel et al. (2018). However, unlike reversal probability and duration, average speed of reversals typically showed only a small decrease over taps (Figure 1C). Leveraging the data produced by the MWT, we investigated the dynamics of the speed of reversal responses in more detail to identify whether parts of the reversal display greater plasticity that is obscured by averaging speed over the entire reversal. We plotted the reversal speed of the first tap and last three taps over the duration of the reversal to observe how the reversal speed profiles changed over the habituation process.

**Figure 1.**
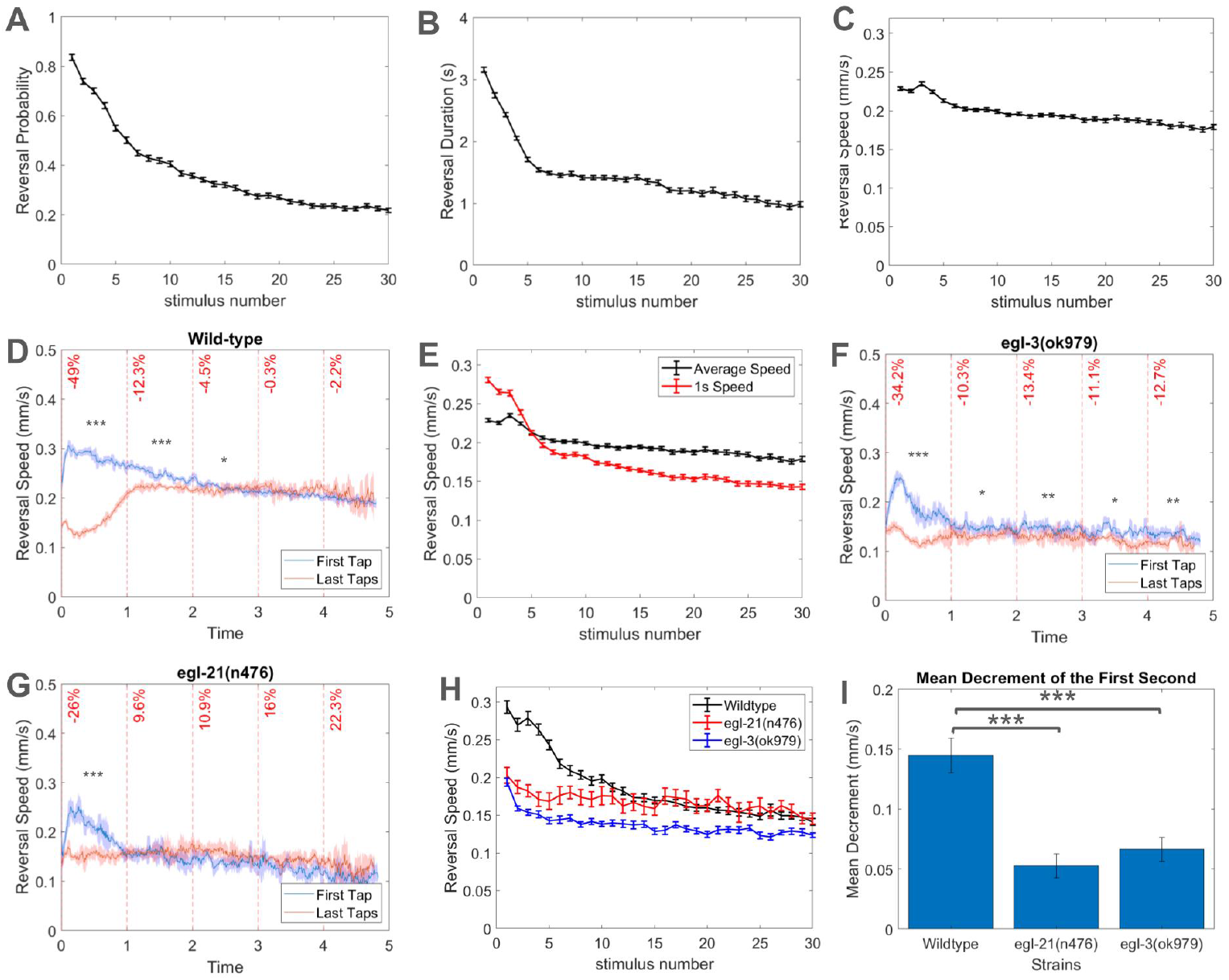
Habituation of speed reversal profiles in *Caenorhabditis elegans*: In all panels, worms were 96 hours post-hatch and were on a lawn of OP50 *E. coli*. Habituation of (A) reversal probability, (B) reversal duration, and (C) reversal speed over 30 taps delivered at a 10s ISI in wild-type *C. elegans*. All error bars are standard errors of the mean. (D) Reversal speed profiles of wild-type worms, with the x-axis showing time after average reversal onset and the y-axis representing reversal speed. The blue line shows the average reversal speed after the first tap and the red line represents the average reversal speed after the final three taps, with a confidence interval of 95%. Percentage decrease (vertical red text) in reversal speed averaged every 1s after reversal onset is overlaid on these graphs. Student t-tests were conducted between the reversal speed of the first and last taps for every 1s (*** = p < 0.0001). (E) Habituation curves comparing the average speed across the entire reversal (black line; mean +/-SEM) with the average speed of the first second of the reversal (red line; mean +/-SEM). (F) Reversal speed profiles over the duration of reversals for *egl-3* mutants with the x-axis showing time after average reversal onset and the y-axis represents reversal speed, the blue line shows the average reversal speed after the first tap and the red line represents the average reversal speed after the final three taps, with a confidence interval of 95%. Percentage decrease (vertical red text) in reversal speed averaged every 1s after reversal onset is overlaid on these graphs. Student t-tests were conducted between the reversal speed of the first and last taps for every 1s (* = p < 0.05, ** = p < 0.001, *** = p < 0.0001). (G) Reversal speed profiles over the duration of reversals for *egl-21* mutants with the x-axis showing time after average reversal onset and the y-axis representing reversal speed. The blue line shows the average reversal speed after the first tap and the red line represents the average reversal speed after the final three taps, with a confidence interval of 95%. Percentage decrease (vertical red text) in reversal speed averaged every 1s after reversal onset is overlaid on these graphs. Student t-tests were conducted between the reversal speed of the first and last taps for every 1s (* = p < 0.05, *** = p < 0.0001). (H) Average reversal speed of the first second for *egl-3* (in blue) and *egl-21* mutants (in red), compared to the wild-type worms they were tested against (in black). (I) The decrease in reversal speed from the first to final taps during the first second after reversal (average reversal speed after first tap - reversal speed after the last three taps) can be seen for wild-type worms, *egl-3* mutants, and *egl-21* mutants, with 95% confidence intervals. Using student t-tests, significant differences of mean speed decrement were seen between mutants and the wild-type worms they were tested against (*** = p < 0.0001).

When wild-type worms were given 30 taps at a 10s interstimulus interval (ISI), the speed of the response to the first tap increased to a peak shortly after reversal onset, and then gradually decreased throughout the rest of the reversal (Figure 1D). In contrast, reversal speed of the final three taps initially dropped to a low level shortly after reversal onset before increasing to the speed similar to the speed 2s after reversal onset of the first tap. The largest change in reversal speed between the first and the final taps was observed within the first second of the reversals. A 49.0% decrease in speed was observed during the first second of reversal, with a significant difference between the reversal speed after the first tap and the last taps (p < 0.0001) and an extremely large effect size of d = 4.43 (Cohen, 1988). From 3s after reversal onset to the end of the reversal, there were no significant differences between the speed of the first and the last taps.

This lack of decrement in reversal speed for much of the reversal explains why graphs of average reversal speed habituation do not show a large response decrement. Averaging the part of the reversal that does habituate with large parts of the reversal that do not habituate over the repeated taps dilutes the observed change in reversal speed. This is most clearly demonstrated when comparing habituation curves for average speed (in black), showing a decrease of 22.4% from tap 1 to tap 30, to habituation curves for average speed of the first second of reversal (in red), which shows a substantially larger behavioral decrease of 49.0% (Figure 1E). The habituation curve for the average speed of the first second of reversal more closely resembles the decrement observed in the probability and duration habituation curves (Figure 1A and 1B) than did the average speed curve (Figure 1C).

These findings highlight the dangers of averaging responses when investigating habituation, as many behavioral shifts in response to stimuli can be subtle and time sensitive. Early studies investigating habituation deficits in patients with schizophrenia were problematic as researchers sometimes compared the average of all responses before training (baseline) with the average of all responses after training (McDiarmid et al., 2017). These analyses would miss any differences in habituation if the rate of habituation is rapid. Our findings extend this idea to averaging over an entire response to a single stimulus. As demonstrated here, it is possible that habituation of a response may occur at a specific moment of the response and analyses that average over the entire behavior may dilute large behavioral shifts with invariant parts of the response.

Previously, our lab showed that neuropeptides, short sequences of amino acids that can act as primary transmitters in *C. elegans*, promote aspects of habituation (*i*.*e*. Ardiel et al., 2017). Utilizing this new measure of reversal speed, we analyzed reversal speed profiles for worms with mutations in genes important for the synthesis and processing of many neuropeptides: *egl-3* (ortholog of proprotein convertase 2; involved in cleaving proneuropeptides) and *egl-21* (ortholog of carboxypeptidase E; involved in removing C-terminal basic residues of peptide intermediates; WormBase, 2024). For *egl-3* (Figure 1F) and *egl-21* mutants (Figure 1G), the reversal speed profiles for the first tap show moderate increases in speed shortly after reversal onset before quickly decreasing to ~0.14 mm/s 1s after reversal onset. The reversal speed of the last taps remained ~0.14 mm/s throughout the reversal. While the greatest difference in reversal speed from the first to the last taps still occurred during the first second of reversal, the difference was much smaller than what was seen for wild-type worms.

Over the course of 30 taps, we compared the reversal speeds of wild-type worms and worms with mutations in *egl-3* and *egl-21* (Figure 1H). Wild-type worms displayed a decrease of 0.145 mm/s during the first second of reversal from the first to the last taps. In contrast, *egl-3* and *egl-21* mutants showed a decrease of 0.066 mm/s and 0.053 mm/s respectively. Within the first second of reversal the mean decrement observed in the mutants was significantly less than the wild-type worms they were tested against (p < 0.0001) (Figure 1I).

While the plasticity of reversal speed was substantially reduced in these mutants, a decrement in reversal speed over 30 taps was still seen, suggesting that there may be additional mechanisms involved in behavioral plasticity of reversal speed. Mutations in *egl-3* and *egl-21* eliminate a large number of neuropeptides, but not all of them, so it is possible that these remaining neuropeptides could also be involved. Further work is required to determine which neuropeptides and receptors mediate plasticity of reversal speed. The major families of neuropeptides include insulin-like peptides, FMRF amide-related peptides, and non-insulin, non-FaRP neuropeptides, most of which are processed by *egl-3* and *egl-21* (Li and Kim 2014). Future behavioral testing of animals with mutations in specific neuropeptide families are required to further elucidate this potential neuropeptide mechanism of reversal speed plasticity.

In summary, we found that speed of tap elicited reversals is quite plastic at the beginning of the reversal in wild-type worms, with some parts of the reversal decreasing from the first to final taps nearly 50%. Immediately after the first tap, reversal speed increased greatly within the first second after reversal onset before gradually slowing down over the remainder of the reversal. For the final taps, reversal speed rapidly dropped, reaching its lowest point within the first second of reversals, before increasing to a speed that was maintained for the rest of the reversal. In contrast to the first second of reversals, the last few seconds of the reversals to the first and the last taps were not significantly different. It is clear that when exposed to taps, worms do not change reversal speed for the entire reversal. Instead, reversal speed is most plastic during the first second of reversal, with little difference in reversal speed later in the reversal. With repeated taps, more habituation can be seen when reversal speed is averaged only for the first second of reversal compared to when it is averaged throughout the reversal. We observed significantly less reversal speed decrement from first to last taps in animals with mutations in neuropeptide processing genes. Overall, this approach to analyzing reversal speed over taps demonstrated that habituation of reversal speed in *C. elegans* occurred primarily in the first second after reversal onset, with this behavioral shift in reversal speed being, at least partially, mediated by neuropeptides.

## Methods

### a) Strain Maintenance and Strains

*C. elegans* were maintained on Nematode Growth Medium (NGM) and seeded with 3-4 drops *Escherichia coli* (OP50) and incubated at 20°C as described in Brenner (1974). 3 adults of each plate were picked to a new plate every 4 days for maintenance.

The strains used in the study have mutations in genes involved in the neuropeptide processing pathway. These strains are VC671 *egl-3*(ok979) and KP2018 *egl-21*(n476).

### b) Behavioural testing

Worms were age synchronized to control for age differences by allowing seven adults (4 days old) to lay eggs for 3-4 hours on OP50 *E. coli* seeded NGM plate (Timbers et al., 2013). The adults were then removed from the plate which were then incubated in a 20°C fridge. After four days, seven more worms from these plates would be transferred OP50 *E. coli* seeded MWT plates to lay eggs for 3-4 hours and removed afterwards. After four more days, these plates were used for behavioural testing

These plates were then placed in the lab’s Multi-Worm Tracker apparatus and recorded with program version 1.2.0.2, which was used to score behavior of 40-120 of *C. elegans* simultaneously (Swierczek et al., 2011). An acclimation period of 600s after placing worms in the apparatus preceded 30 taps administered by a metal solenoid to the side of their petri dish at a 10s interstimulus interval.

For wild-type worms, 122 plates were analyzed, whereas 21 plates were analyzed for *egl-3* mutants and 15 plates for *egl-21* mutants.

### c) Image acquisition of behavior

As described in Swierczek et al. (2011), image acquisition was performed with MWT software (version 1.2.0.2). A Dalsa Falcon 4M30 camera (8 bits; 2352 × 1728 pixels, 31 Hz) and a Rodenstock 60-mm f-number 4.0 Rodagon lens under brightfield illumination (backlit with 4″ × 4.9″ light plate, Schott A08925 with ACE I illuminator) were used to visualize petri dishes in the apparatus. A National Instruments PCIe-1427 CameraLink capture card was run along with the MWT tracking software on personal computers with 3 GHz Intel Core 2 Duo processors and 4 GB of RAM. Choreography analysis software (v. 1.3.0_r1035) was used for tap-response and baseline phenotypic quantification.

### d) Statistics and Calculations

Reversal speeds after the first tap were averaged across plates for every 0.01s, while for the final taps, the final three taps (28-30 taps) were also averaged across plates for every 0.01s. The speeds were normalized by average body length for each plate. These reversal speeds were then plotted according to time since the tap for the first tap and the average of the final three taps. A running mean was applied for the reversal speed profile graphs, with a window of 0.02s before and after the particular timepoint.

Percent decrease between the first and final taps were calculated for every 1s after average reversal onset, the time in which worms started reversing in response to the tap. The percentage was calculated by subtracting the average speed at a certain time after the first tap by the same amount of time to the final three taps averaged, dividing it by the average speed of the final three taps, then multiplying by 100.

For each strain, student t-tests and Cohen’s d were used to evaluate differences in reversal speed between the first tap and final taps every 1s after reversal onset. Cohen’s d was interpreted using descriptors of large effect size=.80, medium effect size=.50, small effect size=.20, as described in Cohen (1988). When comparing strains, student t-tests were used to evaluate differences in decrement of reversal speed during the first second of reversal from the first to the final taps.

## Reagents

| Strain | Genotype | Available from: |
| --- | --- | --- |
| N2 | *Caenorhabditis elegans* | CGC |
| VC671 | *egl-3(ok979)* | CGC |
| KP2018 | *egl-21(n476)* | CGC |

## Acknowledgements

Some strains were provided by the CGC, which is funded by NIH Office of Research Infrastructure Programs (P40 OD010440).

Wormbase

## Author Contributions

Audrey Siu: conceptualization, investigation, methodology, formal analysis, writing - original draft. Nikolas Kokan: conceptualization, methodology, supervision, writing - review editing. Catharine H Rankin: funding acquisition, supervision, writing - review editing.

## Reviewed By

### History

#### Copyright

© 2025 by the authors. This is an open-access article distributed under the terms of the Creative Commons Attribution 4.0 International (CC BY 4.0) License, which permits unrestricted use, distribution, and reproduction in any medium, provided the original author and source are credited.

#### Citation

Siu A, Kokan N, Rankin CH. 2025. Neuropeptides Involved in Elicited Reversal Speed Plasticity in *C. elegans* During Mechanosensory Habituation. microPublication Biology.

